# Distinct cytotoxic cell subsets underlie protective and non-protective immunity to African swine fever virus

**DOI:** 10.64898/2025.12.01.691414

**Authors:** D. Marín-Moraleda, A. Tort-Miró, E. Ezcurra, S. Montaner-Tarbes, J. Muñoz-Basagoiti, E. Coatu, M.J. Navas, M. Muñoz, P. Monleon, J. González-Oliver, L. Pailler-García, S. Pina-Pedrero, F. Accensi, F. Rodríguez, J. Argilaguet

**Author notes:** These authors contributed equally to this work (D. Marín-Moraleda, A. Tort-Miró).

## Abstract

Limited understanding of African swine fever (ASF) immunity remains a major barrier to the rational development of safe and effective vaccines. While antibody-mediated protection is still poorly defined, growing evidence highlights a central role for cellular immunity. In particular, cytotoxic cells have emerged as key components to control ASF virus (ASFV) infection. However, the contribution of individual cytotoxic subsets across different virological and immunological contexts is not well characterised. Here, we investigated cytotoxic responses during BA71ΔCD2 live attenuated vaccine (LAV)-induced protection and during late-stage lethal ASFV infection, and demonstrated the involvement of different cytotoxic subsets in each scenario. Early increases in perforin-producing CD8αβ^+^ T cells in blood after immunisation coincided with the onset of protection. At later time points, elevated levels of these cells after *in vitro* ASFV-specific stimulation correlated with survival to lethal challenge, supporting their central role in protective immunity. Additional correlates of protection during recall responses included CD4^+^CD8αβ^+^ cytotoxic T cells, IFN*γ*-producing cells, and ASFV-specific antibodies, illustrating the multifactorial nature of immunity to ASF. In contrast, pigs with acute ASF exhibited a distinct cytotoxic profile characterised by broad increases across multiple perforin-producing subsets. Although all of them showed reduced susceptibility to ASFV-induced lymphopenia, only perforin-producing NK and *γ*δ T cells correlated with viremia, suggesting their active involvement during late disease. Together, these findings advance our understanding of cytotoxic responses to ASFV and identify cytotoxic T cells, alongside other immune components, as potential correlates of protection that may guide future vaccine development.

## INTRODUCTION

The complexity of immune responses to African swine fever (ASF) remains one of the major obstacles to developing safe and effective vaccines against this devastating disease. The causative agent, African swine fever virus (ASFV), continues to spread across Africa, Asia and Europe through the infection of domestic pigs and wild boars and causing substantial economic losses to the global swine industry^1^. Live attenuated vaccines (LAVs) induce strong protective immunity^2–4^, and, notably, two LAVs were approved in Vietnam in 2022^5^. However, biosafety concerns limit their wider deployment^6–9,^ highlighting the need for safer subunit vaccines. In this line, progress has been made in identifying protective antigens and delivery platforms^10–12^, but vaccine development continues to be hindered by the complexity of ASFV, a large DNA virus encoding over 150 proteins^13^ with multiple circulating genotypes^14^. Moreover, the immune mechanisms underlying both protective and pathological responses remain poorly defined, further limiting rational vaccine design^2,15^.

The pathogenesis of acute ASF is a direct consequence of the virus’ capacity to modulate host immune responses at multiple levels. ASFV functionally manipulates and kills infected monocytes and macrophages by interfering with innate and apoptotic signalling pathways^16,17^. As a result, infected cells fail to mount an antiviral state or efficiently prime neighbouring immune cells to develop an effective protective immunity^18^. In addition, ASFV induces severe leukopenia through the bystander death of uninfected cells, mediated by viral proteins and dysregulated production of inflammatory cytokines^19,20^. Combined, these immune evasion strategies lead to rapid immunosuppression, preventing the host from developing a timely protective immunity and resulting in a fatal outcome. This pathological profile contrasts with the non-aggressive, self-limiting infections induced by LAVs or naturally attenuated strains. While mechanisms of attenuation are diverse^4^, all these ASFV strains replicate while causing limited immune disruption and therefore allow the host to mount effective adaptive responses. Consequently, attenuated strains have been widely used as experimental tools to characterize protective immune mechanisms against ASFV^21,22^.

Host immune responses to ASFV are complex and highly context-dependent. Although antibodies play a relevant but limited role in protection^23–25^, complementary cellular immunity is necessary for controlling infection^26^. Increasing evidence indicates that cytotoxic responses are central in protective immune responses. For instance, depletion of CD8α^+^ cells in pigs immunised with a naturally attenuated strain significantly impaired protection^27^, and most cytotoxic subsets express CD8α^28^. Moreover, memory cytotoxic T cells are induced in pigs immunised with various LAVs or attenuated ASFV strains^26,29–33^. In some contexts, these responses also involve cytotoxic *γ*δ T and NK cells^31,34,35^, highlighting the multifaceted nature of cytotoxic immunity. Additionally, post-challenge analyses further indicate that both CD8β^+^ T cells and NK cells may contribute to virus control and clearance^27,36^. Adding complexity, acute ASF is characterised by early reductions in cytotoxic subsets^37–39^, followed by increased cytotoxic activity during advanced disease^26,40–42^.

In this study, we aimed to clarify the role of cytotoxic responses across distinct virological and immunological contexts following LAV immunization or ASFV infection. Using the BA71ΔCD2 LAV prototype, which induces a dose-dependent protection^31^, we show that CD4^−^CD8αβ^+^ T cells are a critical component of protection. BA71ΔCD2 immunization induced an early, gradual increase in circulating CD4^−^CD8αβ^+^ T cells that correlated with the onset of protection at 12 days post-vaccination. Additionally, a comprehensive analysis of memory responses revealed that both cellular and humoral components correlate with cross-protection against the Georgia2007/1 challenge, with CD4^−^CD8αβ^+^ T cells showing the strongest association with infection control. In contrast, pigs with acute ASF displayed broad increases in multiple cytotoxic subsets, including CD4^−^CD8αβ^+^ T cells, CD4^+^CD8αβ^+^ T cells, CD8αα^+^ *γ*δ T cells, and NK cells, which were less susceptible to ASFV-induced lymphopenia than their non-cytotoxic counterparts. Importantly, among these expanded subsets, only perforin-producing NK cells and CD8αα^+^ *γ*δ T cells correlated with viremia. Overall, this study provides new insights into the role of cytotoxic lymphocytes in ASF immunity and identifies cytotoxic T cells, together with complementary immune components, as key correlates of protection, contributing to the rational design of ASF vaccine strategies.

## METHODS

### Ethics statement

Animal care and experimental procedures were carried out in accordance with Good Experimental Practice guidelines and were approved by the Ethics Committee on Animal Experimentation of the *Generalitat de Catalunya* (protocols codes 12433, 12393, 12648 and 11947). Protocol codes corresponding to the cryopreserved samples from published studies are provided in the respective original publications^40,43^. All *in vivo* experiments were conducted in biosafety level 3 facilities at the Centre de Recerca en Sanitat Animal (IRTA-CReSA, Barcelona).

### Samples from animal infections

Samples from domestic pigs were obtained from several experiments. PBMCs from BA71ΔCD2-vaccinated pigs originated from two previously published experiments^31,40,43^. For analyses of pigs undergoing acute ASF, samples were derived from either cryopreserved or fresh PBMCs collected from unvaccinated animals infected with the Georgia2007/1 strain. Cryopreserved samples were obtained from three independent studies: two involving intranasally challenged pigs (10^5^ HAU; n= 11) and one involving pigs (n= 5) infected through an in-contact transmission model, as previously described^31^. In addition, a dedicated *in vivo* experiment was conducted to quantify absolute counts of cytotoxic subsets in fresh PBMCs from intranasally infected pigs (10^5^ HAU; n= 5). In all cases, six-to eight-week-old domestic pigs underwent a seven-day acclimatation period and were fed *ad libitum* throughout the experiment. Serum and EDTA-blood samples were collected at the indicated time points. Animal health status was monitored daily according to standardised welfare guidelines. Clinical signs were evaluated following the criteria described by Galindo-Cardiel et al.^44^, assessing behaviour, body condition, presence of cyanosis, and digestive and respiratory signs. Each parameter was scored on a scale from 0 to 3 (0 = normal; 1 = mild; 2 = moderate; 3 = severe). Animals that reached humane endpoints were humanely euthanised.

### Viruses

BA71ΔCD2 is a LAV prototype obtained by deletion of the CD2v gene (EP402R) from the parental virulent BA71 ASFV strain^45^. BA71ΔCD2 was expanded in the established COS-1 cell line (ATCC) and titrated by immunoperoxidase monolayer assay (IPMA) as previously described^45^. The virulent Georgia2007/1 ASFV strain was kindly provided by Dr. Linda Dixon (WOAH reference laboratory, Pirbright Institute, UK), expanded in PAMs, and titrated by hemadsorption assay as previously described^46^. Titres were expressed as TCID_50_/mL and HAU_50_/mL, respectively, according to the Reed–Müench method^47^. TCID_50_ values were converted into plaque-forming units (pfu) applying a Poisson distribution (TCID_50_/mL = 0.7 pfu/mL).

### Quantitative PCR for the detection of ASFV

Viral genomic DNA was extracted from blood or sera by IndiMag® Pathogen Kit (INDICAL Bioscience, Leipzig, Germany) in a semi-automated manner using the KingFisher System (Thermo Fisher Scientific, Waltham, MA, USA). Viral titres were assessed by SYBR Green qPCR targeting the ASFV PK gene as previously described^45^. Ten-fold serial dilutions of a plasmid encoding the PK gene were used as standard template to determine the sensitivity of the assay.

### Flow cytometry

To assess cytotoxic recall responses, cells were stimulated with either BA71ΔCD2 or Georgia2007/1 at a MOI of 0.2 for 36 hours at 37ºC. RPMI medium and phorbol myristate acetate (PMA, 5 ng/mL) plus ionomycin (500 ng/mL) were included as negative and positive controls, respectively. Analyses of cytotoxic subsets in PBMCs from naive or acutely infected pigs were performed *ex vivo* without previous stimulation. In all cases, 10^6^ fresh or cryopreserved PBMCs were used per condition in U-bottom 96-well plates, and staining of cells for flow cytometry was performed as previously described^31^. Briefly, PBMCs were stained with LIVE/DEAD Fixable Red Dead Cell Stain Kit (ThermoFisher Scientific) following manufacturer’s instructions. Blockage of Fc receptors was performed for 15 min on ice prior to antibody staining. Staining of cells to analyse frequencies of cytotoxic subsets was perfromed with an antibody mix containing mouse anti-pig CD3e (PE-Cy7, Clone BB23-8E6-8C8, BD Biosciences), mouse anti-pig CD4α (PercP-Cy5.5, Clone 74-12-4, BD Biosciences), rat anti-pig *γ*δTCR (APC, Clone MAC320, BD Biosciences), anti-pig CD8α (FITC, Clone 76-2-11, BD Biosciences) and mouse anti-pig CD8β (R-PE, Clone PPT23, Bio-Rad Laboratories). After extracellular staining, cells were fixed and permeabilised with the BD Cytofix/Cytoperm Kit (BD Biosciences) and intracellularly stained with mouse anti-human/pig perforin (eFluor450, Clone δG9, eBioscience). To immunophenotype cytotoxic T cells, the anti-CD8α and anti-*γ*δTCR antibodies were replaced with the mouse anti-pig CD45RA (FITC, Clone MIL13, Bio-Rad Laboratories) and the rat anti-human CD197 (CCR7) (APC, Clone 3D12, Invitrogen). Samples were acquired on a BD FACSAria IIu flow cytometer (BD Biosciences), and data was analysed using FlowJo v10.7.1 software (Tree Star Inc).

### Microfluidic quantitative PCR assay

PBMCs from BA71ΔCD2-vaccinated pigs were seeded at 10^7^ cells/well in 6-well flat-bottom plates and overnight stimulated with either BA71ΔCD2 or Georgia2007/1 strains (both at MOI 0.2). Total RNA was extracted using the RNeasy Mini Kit (Qiagen), and the concentration and integrity were measured using the Qubit RNA BR Assay Kit (ThermoFisher Scientific) and the Agilent Bioanalyzer, respectively. cDNA was generated from 160 ng of total RNA using the PrimeScript RT reagent Kit (Takara, Japan). The microfluidic qPCR protocol and the primers used are detailed in Bosch-Camos et al.^31^. Briefly, gene expression was quantified in duplicate using the 96.96 Dynamic Array integrated fluidic circuit of the Biomark HD system (Fluidigm Corporation). Data was analysed using the Fluidigm Real-Time PCR Analysis software v4.1.3 and the DAG Expression software v1.0.5.6^48^. Relative quantification of target gene expression was performed by normalising to the average expression of three reference genes (YWHAZ, RPL4 and GAPDH) using the 2^−ΔΔCt^ method and expressed as fold change (FC) relative to the unstimulated control samples. Log_2_-transformed FC values were used for downstream analyses. Z-scores were calculated from individual log_2_FC values for heatmap visualisation, whereas mean log_2_FC values per animal were used for correlation analyses.

### Statistical analyses

Statistical analyses were performed using GraphPad Prism version 10.6.1 (GraphPad Software). The specific statistical tests applied to each dataset are indicated in the corresponding figure legends, and statistical significance was set at: ns p > 0.05, * p ≤ 0.05, ** p ≤ 0.01, *** p ≤ 0.001, **** p ≤ 0.0001. Graphs were generated using GraphPad Prism and BioRender.com.

## RESULTS

### Early increase of cytotoxic CD8αβ^+^ T cells in BA71ΔCD2-vaccinated pigs correlates with protection

To assess the contribution of cytotoxic lymphocytes to the development of protective immunity against African swine fever (ASF), we used the live attenuated vaccine (LAV) prototype BA71ΔCD2 as a model^31^. We first analysed cryopreserved samples from a previously published study that characterised the onset of protective immunity induced by intranasal vaccination with an optimal dose (10^6^ pfu/pig) of BA71ΔCD2^40^. In that study, pigs were challenged intranasally at 3, 7 or 12 days post-vaccination (3dpv, 7dpv and 12dpv groups), while unvaccinated animals served as controls (**Figure 1A**). As reported, pigs challenged at 3 or 7dpv showed only a slight delay in disease progression, whereas animals challenged at 12dpv were able to control infection^40^. Importantly, blood transcriptomic profiling revealed that the onset of protective immunity at 12dpv was associated with a strong cytotoxic signature^40^. Here, we aimed to determine which cytotoxic cell subsets contribute to this early BA71ΔCD2-induced protection. Frozen peripheral blood mononuclear cells (PBMCs) collected immediately before challenge (0 dpc) were analysed by flow cytometry to quantify perforin-producing CD4^−^CD8αβ^+^ T cells, CD4^+^CD8αβ^+^ T cells, CD8αα^+^ *γ*δ T cells and NK cells (**Figure 1A** and **Figure S1**). Only CD4^−^CD8αβ^+^ T cells were significantly increased in the protected 12dpv group (**Figure 1B**). All other cytotoxic subsets showed similar levels across groups, indicating no association with vaccine-mediated protection.

**Figure 1.**
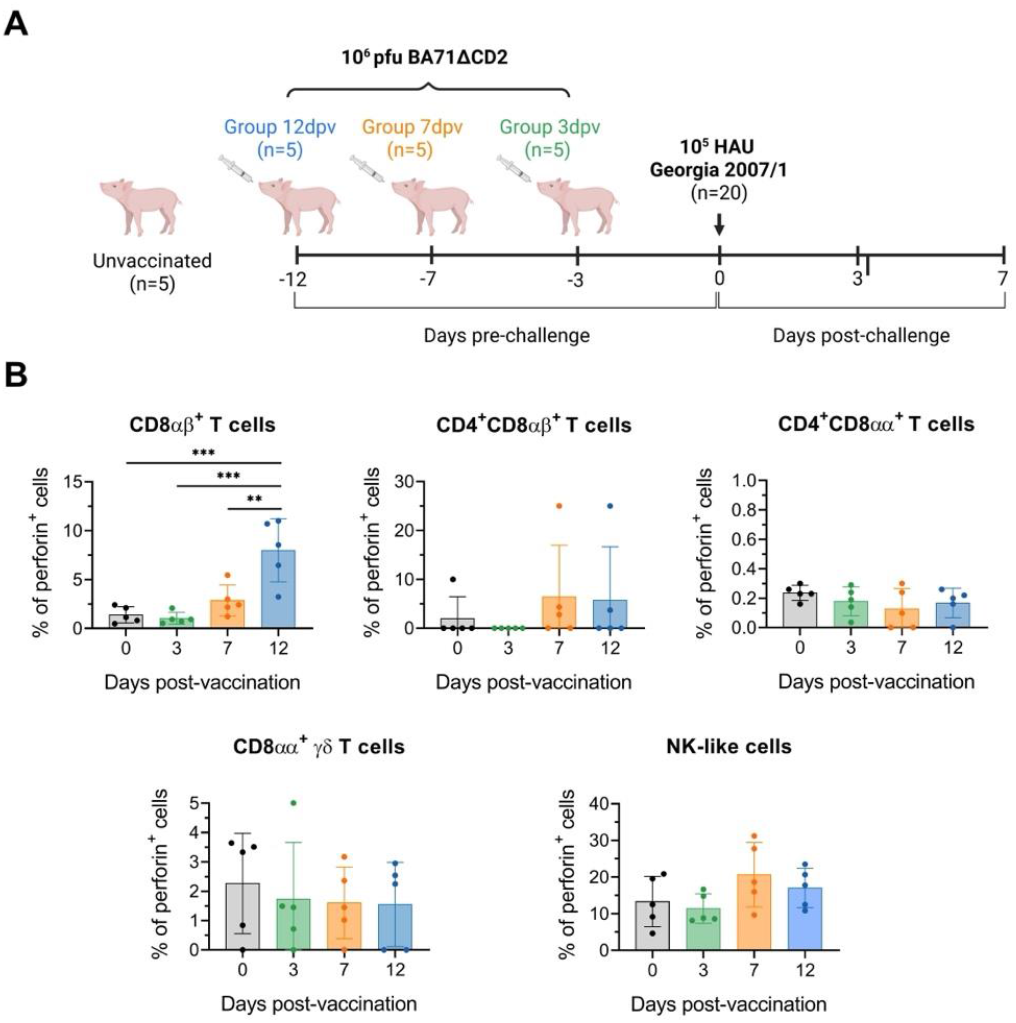
Expansion of blood cytotoxic CD8αβ^+^ T cells 12 days after BA71ΔCD2 vaccination. **A)** Schematic representation of the experimental design from Marín-Moraleda et al.^40^. **B)** Percentages of perforin-producing blood cells (within the parental populations) in four experimental groups: unvaccinated (day 0; black), 3dpv (day 3 post-vaccination; green), 7dpv (day 7 post-vaccination; orange), and 12dpv (day 12 post-vaccination; blue). Analyses were performed on PBMCs without *in vitro* stimulation by flow cytometry. Statistical significance was determined by one-way ANOVA followed by Tukey’s multiple comparisons test (** p ≤ 0.01; *** p ≤ 0.001).

We next performed correlation analyses between cytotoxic cell frequencies and protection. For each pig from the three vaccinated groups, we calculated the area under the curve (AUC) for rectal temperatures, clinical scores and viremia measured at different days post-challenge (**Figure S2**) and compared these values with pre-challenge cytotoxic cell frequencies (**Figure 1B**). Strikingly, only perforin-producing CD4^−^CD8αβ^+^ T cell correlated with reduced fever, milder clinical scores and lower viremia (**Figure 2**). The other cytotoxic subsets did not correlate with any clinical parameters.

**Figure 2.**
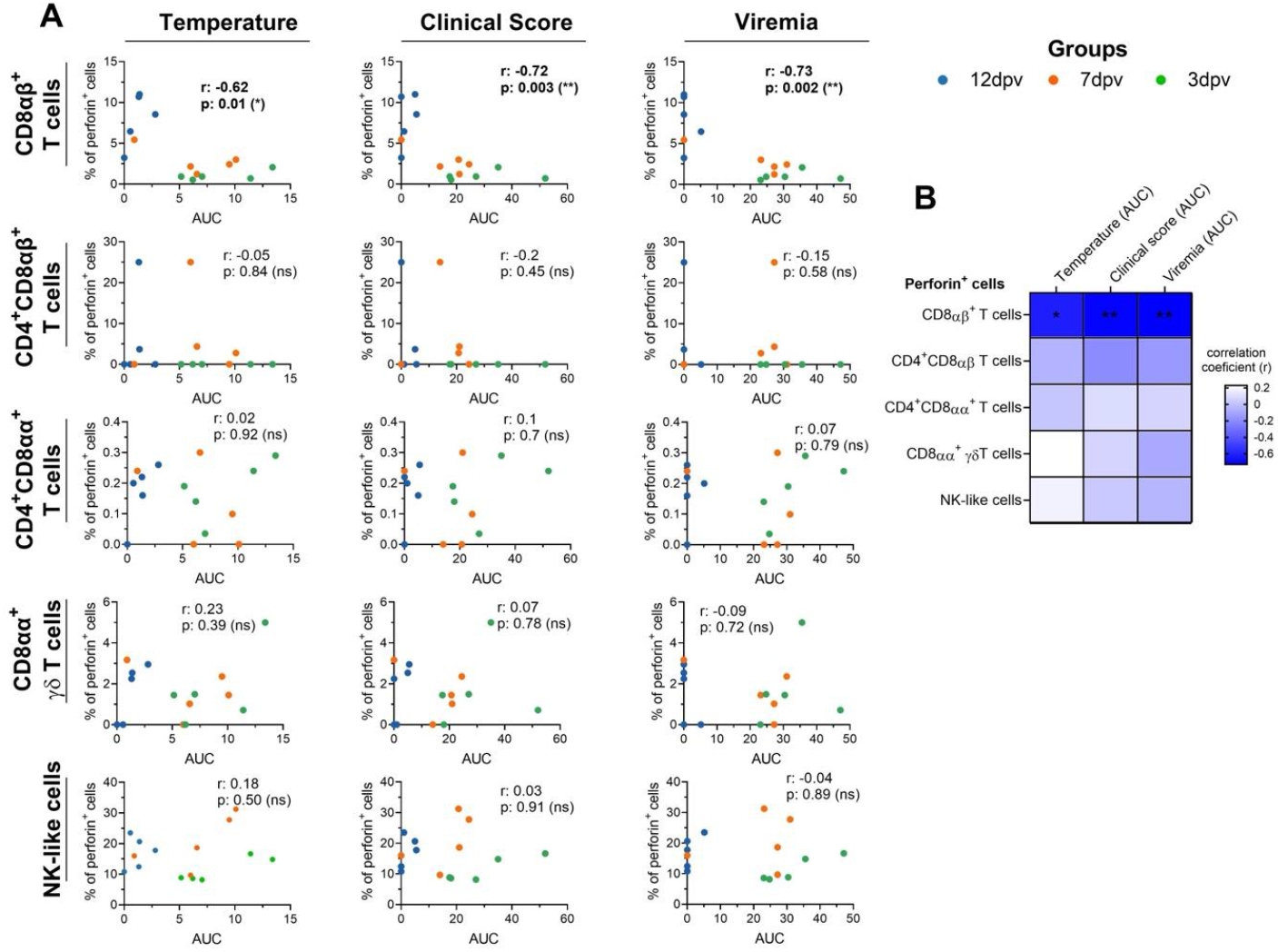
BA71ΔCD2-induced CD8αβ^+^ cytotoxic T cells in blood correlate with disease control. **A**) Correlation between frequencies of perforin-producing cells before challenge and normalised AUC values of rectal temperature, clinical score or viremia after challenge. Dot colours represent vaccinated groups: 3dpv (green), 7dpv (orange), and 12dpv (blue). **B)** Heatmap showing correlation coefficients (r) and significance values for all relationships between ASFV-specific immune parameters and ASF disease control. Correlations were assessed using Pearson’s or Spearman’s test (p > 0.05 ns; * p ≤ 0.05; ** p ≤ 0.01). Statistically significant correlations are highlighted in bold.

### BA71ΔCD2-induced memory cytotoxic CD8αβ^+^ T cells are a key component of multifactorial protective immunity

In addition to cytotoxic T cells, BA71ΔCD2 vaccination induces virus-specific antibodies and polyfunctional Th1 memory responses, including IFN*γ*-secretion that rapidly activates innate antiviral immunity during recall responses^31^. We therefore sought to determine the relative importance of cytotoxic cells among these vaccine-induced immune components. With this aim, we used samples from a previous study in which pigs received a suboptimal intranasal dose of BA71ΔCD2 alone or in combination with two adjuvant prototypes (HI-Ro and Frac-Ro)^43^ (**Figure 3A**). The adjuvants impaired vaccine efficacy, resulting in a wide range of outcomes, from acute lethal ASF comparable to unvaccinated controls, to partial protection, to complete survival^43^. This heterogeneity enabled identification of correlates of protection.

**Figure 3.**
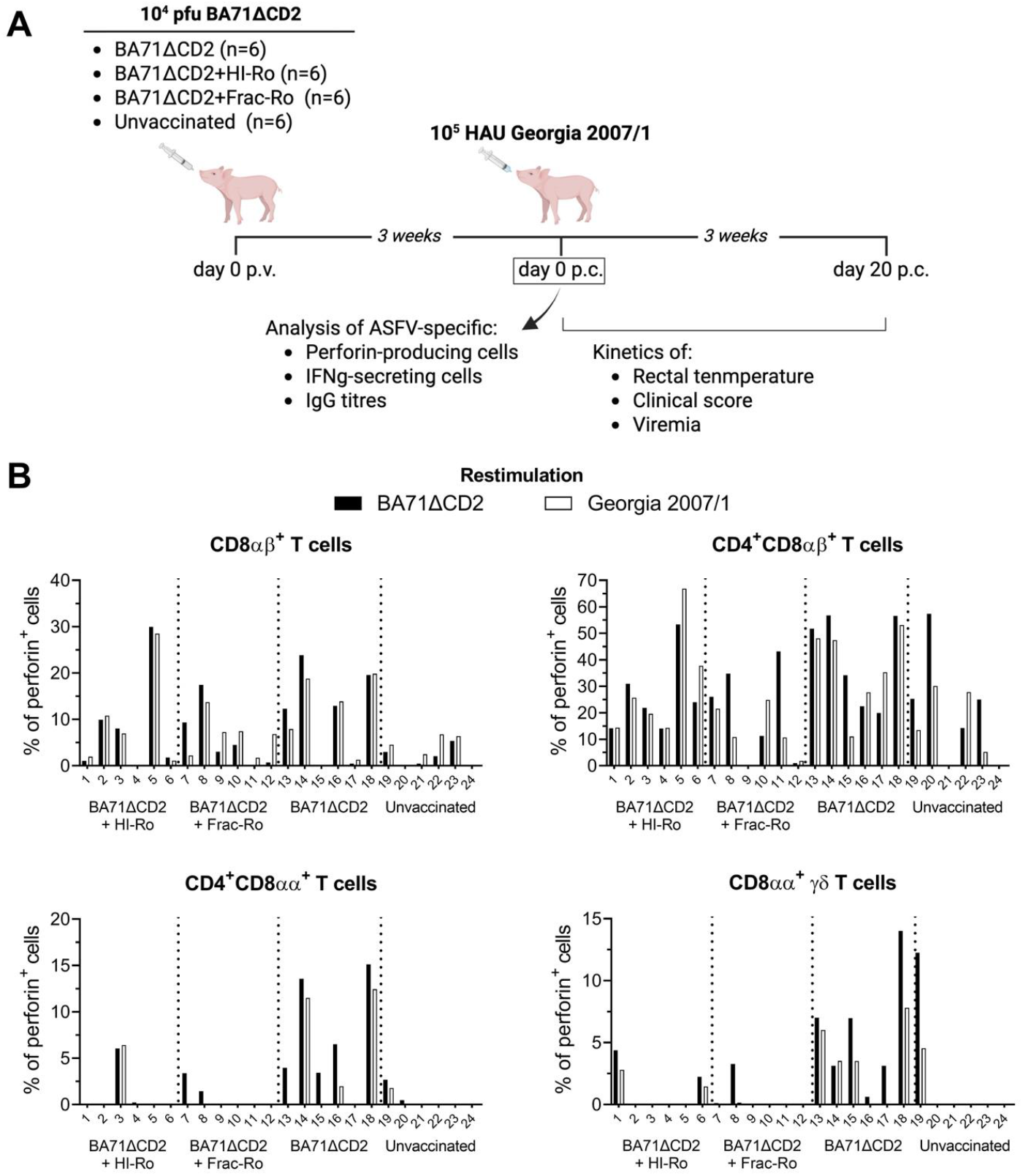
ASFV-specific cytotoxic T-cell responses in PBMCs from pigs vaccinated with BA71ΔCD2 alone or with adjuvant prototypes HI-Ro or Frac-Ro. **A)** Schematic representation of the experimental design from Tort-Miro et al.^43^. **B)** Percentages of perforin-producing cell subsets (within the parental populations) in PBMCs at day 0 post-challenge following *in vitro* stimulation with BA71ΔCD2 (black bars) or Georgia2007/1 (white bars), measured by flow cytometry.

PBMCs collected three weeks after vaccination were stimulated *in vitro* with either the virulent Georgia2007/01 strain or BA71ΔCD2 to assess recall responses (**Figure 3A)**. Perforin-producing cells were quantified by flow cytometry and IFN*γ*-secreting cells by ELISpot (**Figure 3B and Figure S3A**). Although responses varied among animals, the strongest cytotoxic responses were observed within the CD4^+^CD8αβ^+^ population; however, this subset also showed high background levels in some unvaccinated pigs, indicating lower specificity (**Figure 3B**). Correlation analyses using AUCs for rectal temperature, clinical score and viremia (**Figure S3B**) revealed that IFN*γ*-secreting cells and both CD4^+^ and CD4^−^ CD8αβ^+^ cytotoxic T-cell subsets were significantly associated with protection (**Figure 4 and Figure S4**). In contrast, perforin-producing CD4^+^CD8αα^+^ and CD8αα^+^ *γ*δ T cells did not correlate with any clinical parameter. Consistent with our earlier findings (**Figure 2**), the strongest correlation was obtained between protection and cytotoxic CD8αβ^+^ T cells when analysed against clinical scores, highlighting their central role in ASF control (**Figure 4 and 6C**). To further characterize this cell subset, we assessed the expression of CD45RA and CCR7 to distinguish naïve (CD45RA^+^CCR7^+^), central memory T cells (TCM) (CD45RA^−^CCR7^+^), effector memory T cells (TEM) (CD45RA^−^CCR7^−^) and differentiated effector T cells (TDE) (CD45RA^+^CCR7^−^)^49^. Protected pigs showed high frequencies of perforin-producing CD8β^+^ T cells after stimulation, distributed between TEM and TDE phenotypes (**Figure S5**).

**Figure 4.**
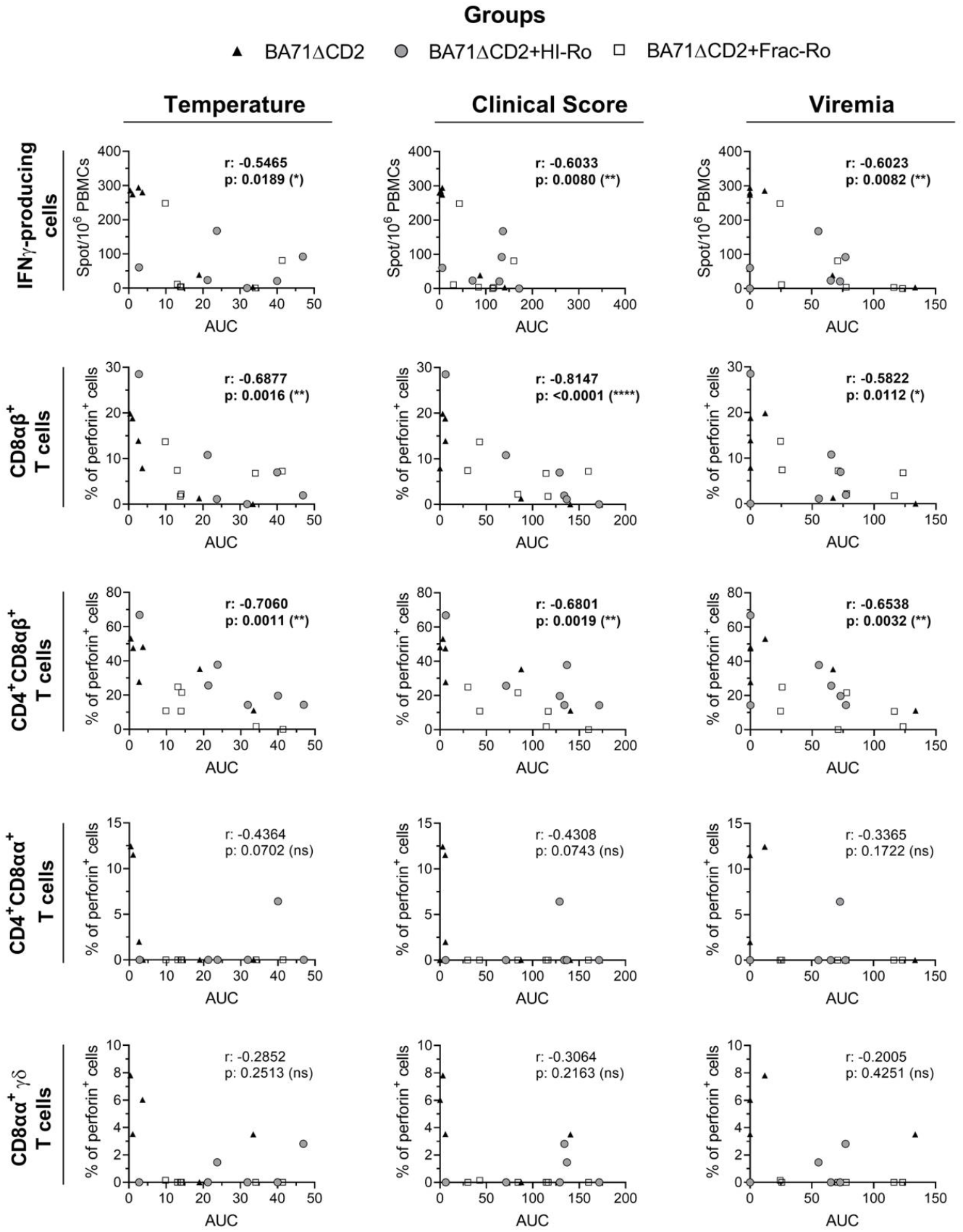
BA71ΔCD2-induced ASFV-specific cellular responses correlate with protection. Frequencies of IFN*γ*-secreting and perforin-producing PBMCs (within the parental populations) before challenge were quantified after stimulation with Georgia2007/1 and correlated with normalised AUC values of rectal temperature, clinical scores, or viremia. Black triangles: BA71ΔCD2 alone; grey circles: BA71ΔCD2+HI-Ro; open squares: BA71ΔCD2+Frac-Ro. Correlations were assessed using Pearson’s or Spearman’s test as appropriate (p > 0.05 ns; * p ≤ 0.05; ** p ≤ 0.01; **** p ≤ 0.0001). Significant correlations are highlighted in bold.

We next contextualised the contribution of cytotoxic CD8αβ^+^ T cells by analysing additional immune components induced by BA71ΔCD2 vaccination. First, we quantified IFN*γ*-driven innate responses through microfluidic qPCR analysis of 34 innate immune-related genes following stimulation of PBMCs with either Georgia2007/1 or BA71ΔCD2. Expression profiles varied across animals, with some showing pronounced upregulation and others resembling unvaccinated controls (**Figures 5 and Figure S6**). Using the mean fold-change between stimulated and unstimulated cells of all genes as a proxy for innate recall activation, we found a significant but modest correlation with protection regardless of the stimulating virus (**Figure 6A**). Notably, innate activation strongly correlated with IFN*γ*-secreting PBMCs (**Figure S7**), supporting the link between inflammatory recall responses and vaccine-induced memory T cells^31^. Finally, ASFV-specific IgG antibodies (**Figure S8**) also correlated significantly with protection (**Figure 6B**), confirming that BA71ΔCD2-induced immunity is multifactorial and includes several potential correlates of protection (**Figure 6C and Figure S9**).

**Figure 5.**
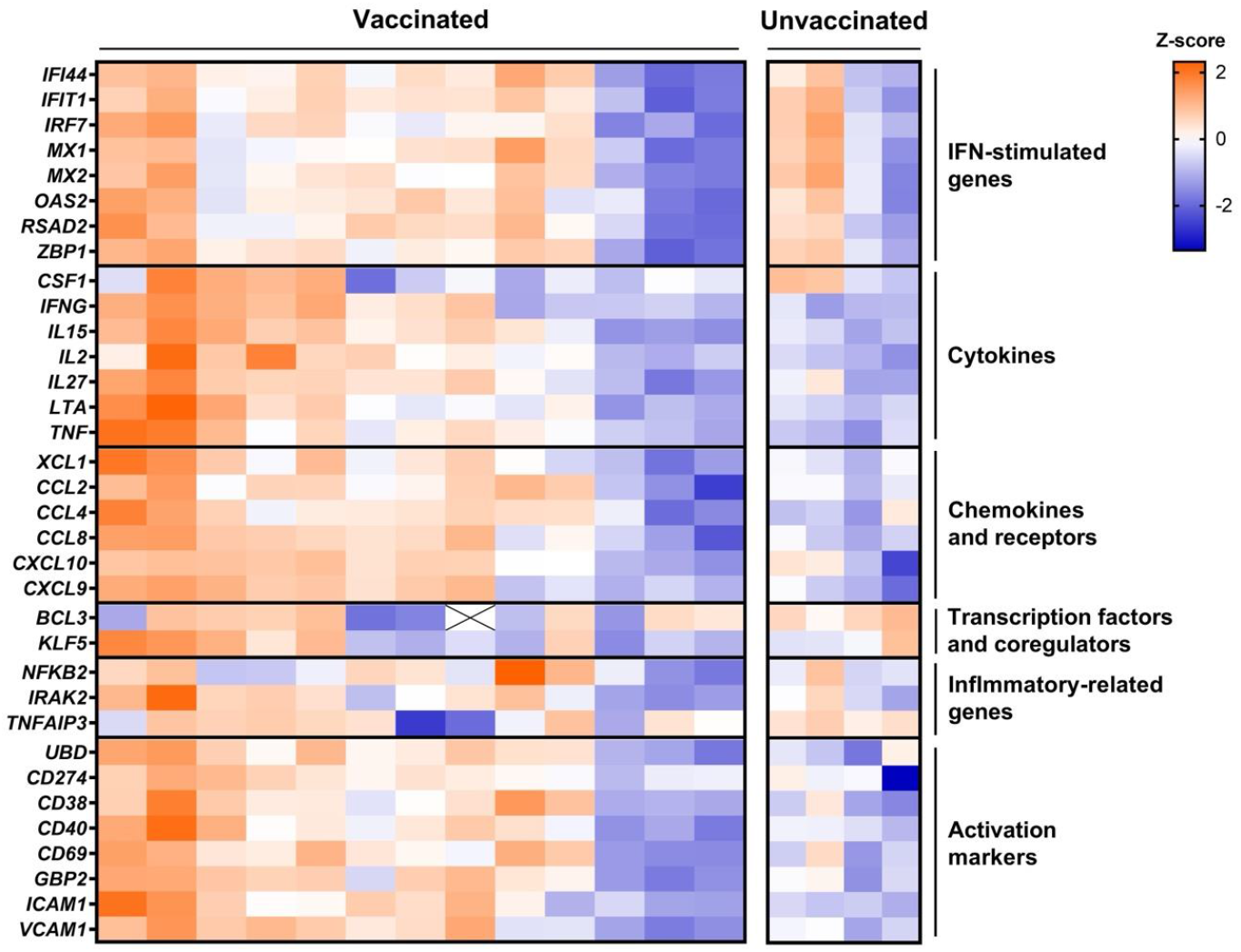
Innate immune transcriptomic signatures during the recall response. PBMCs collected at day 0 post-challenge were stimulated with ASFV Georgia2007/1. Expression levels of innate immune-related genes were quantified by microfluidic qPCR assay. Heatmap shows z-score normalised log_2_ fold changes (log_2_FC) between stimulated and unstimulated cells.

**Figure 6.**
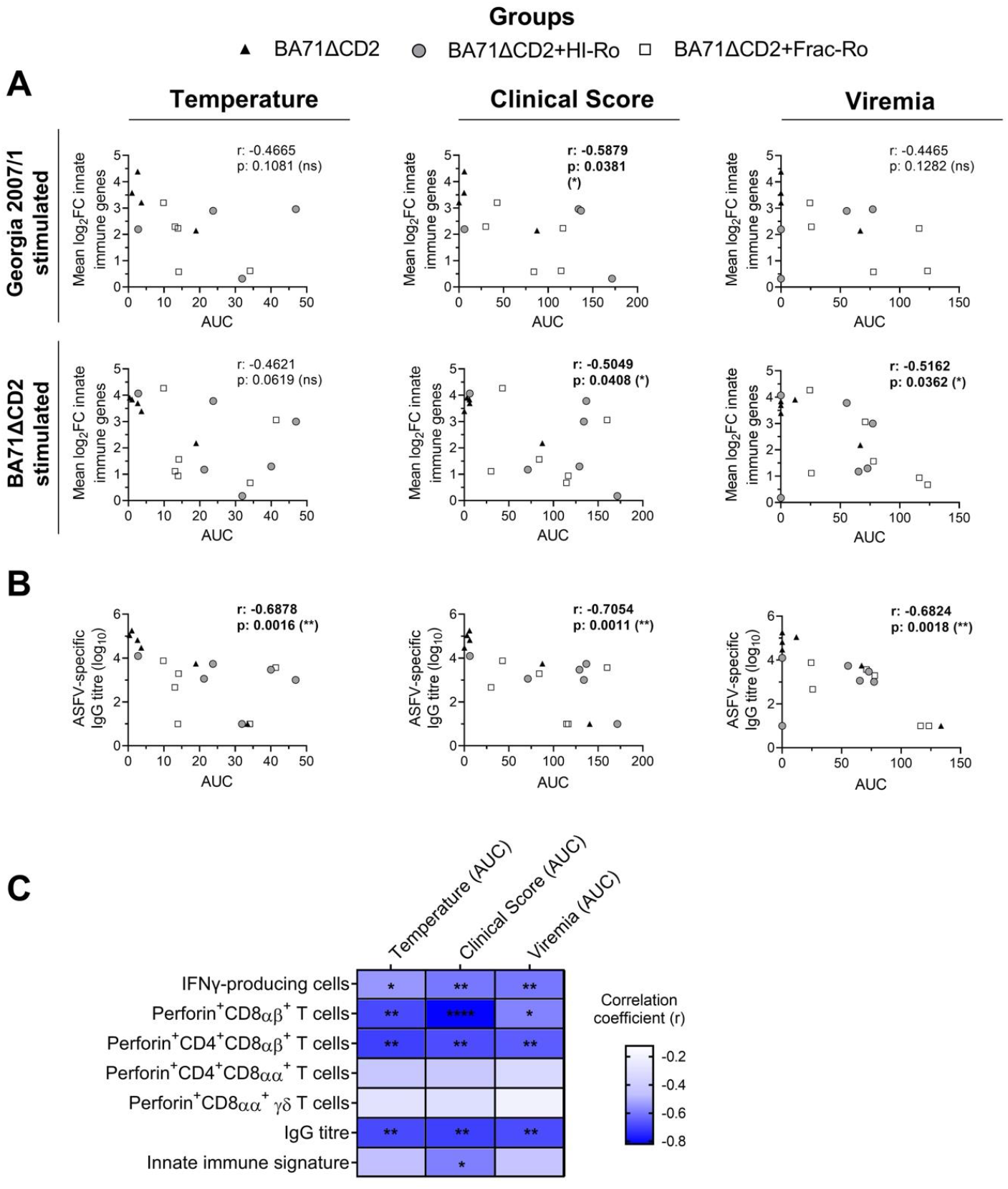
Multiple immune correlates of protection linked to ASF disease control. Correlation analyses between normalised AUC values for rectal temperature, clinical scores, or viremia and **(A)** mean of the log_2_ fold change (log_2_FC) of innate immune gene expression or **(B)** ASFV-specific IgG titres. Mean log_2_FC values were calculated from gene expression in Georgia2007/1-stimulated versus unstimulated PBMCs. Symbols: black triangles (BA71ΔCD2 alone), grey circles (BA71ΔCD2+HI-Ro), open squares (BA71ΔCD2+Frac-Ro). **C)** Heatmap summarising all correlations between ASFV-specific immune variables and disease control. Correlations were assessed using Pearson’s or Spearman’s test (p > 0.05 ns; * p ≤ 0.05; ** p ≤ 0.01; **** p ≤ 0.0001). Significant correlations are highlighted in bold.

### Ongoing acute ASF is associated with broad cytotoxic responses

Cytotoxic activity is also known to increase during acute ASF^26,40,41^. We therefore characterised cytotoxic responses during acute infection and compared them with the protective responses described above. Using cryopreserved PBMCs from the experiment shown in **Figure 1A**, we quantified perforin-producing subsets at 0 and 7 days post-challenge with Georgia2007/01. All cytotoxic subsets were significantly increased at 7 days post-infection in unvaccinated pigs (**Figure 7**). In vaccinated animals, the magnitude of this broad cytotoxic response decreased proportionally with the level of vaccine-induced protection, i.e., pigs challenged at 3 or 7dpv (both unprotected groups) showed moderate cytotoxic responses, whereas fully protected 12dpv pigs maintained basal levels (**Figure 7**). To validate these findings in a larger cohort, we analysed cytotoxic subset frequencies in cryopreserved PBMCs from pigs challenged either intranasally or by contact with Georgia2007/1. Animals euthanised during late-stage acute disease showed robust increase in cytotoxic subsets compared with uninfected controls (**Figure 8A**). Correlation analysis shows that only innate cytotoxic NK-like and CD8αα^+^ *γ*δ T cells correlated with viremia at the time of euthanasia (**Figure 8B and C**), whereas none of the adaptive CD8αβ T-cell subsets showed significant associations.

**Figure 7.**
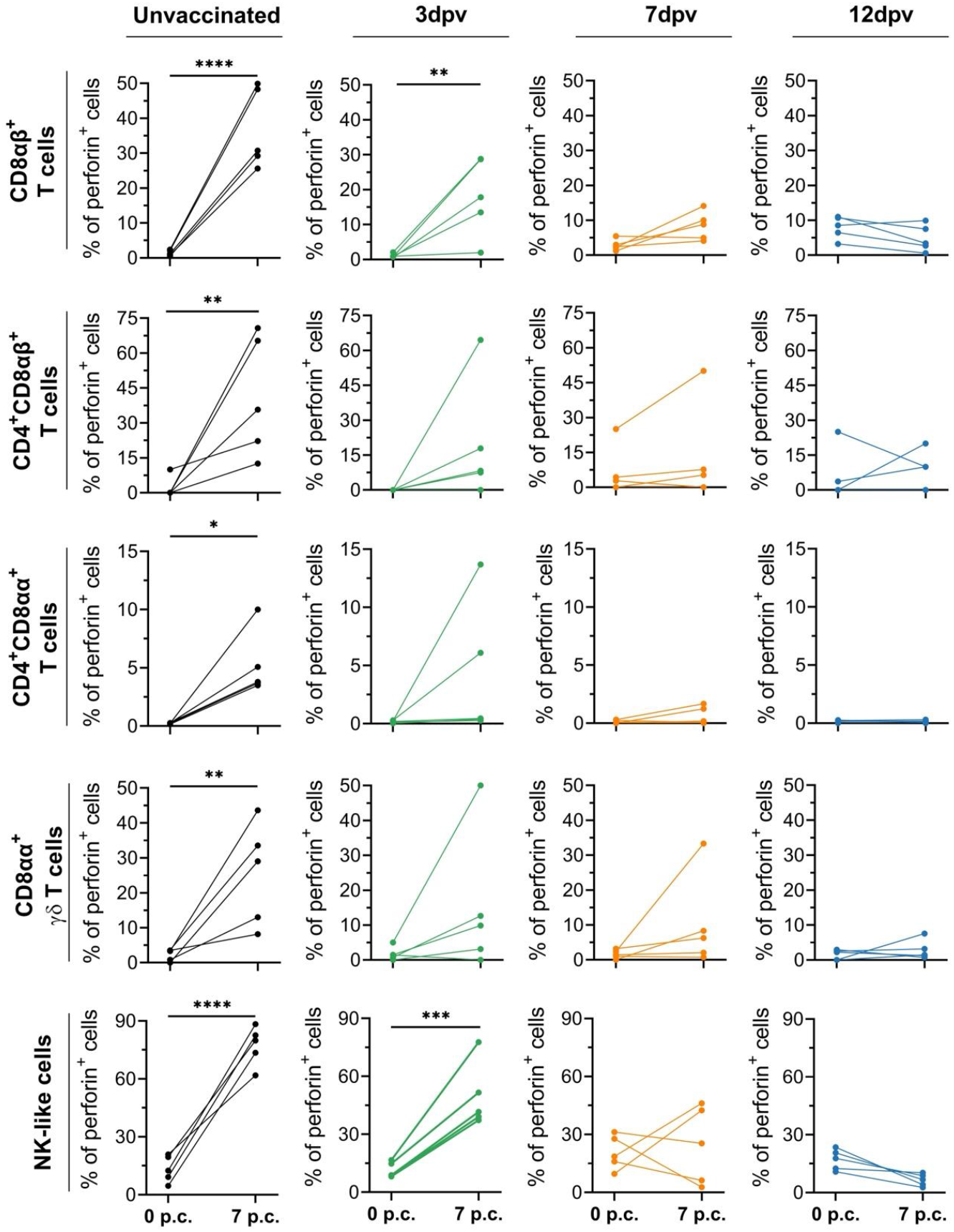
Broad increase of cytotoxic cells in blood during acute ASF. Percentages of perforin-producing blood cells (within the parental populations) at days 0 and 7 post challenge (p.c.) with Georgia2007/1 from unvaccinated pigs (black), and vaccinated animals challenged at day 3 (green), 7 (orange) or 12 (blue) post-vaccination^40^. Analyses were performed on cryopreserved PBMCs analysed by flow cytometry without prior *in vitro* stimulation. Statistical significance was determined by two-way ANOVA followed by Šidák’s multiple comparisons test (* p ≤ 0.05; ** p ≤ 0.01; *** p ≤ 0.001; **** p ≤ 0.0001).

**Figure 8.**
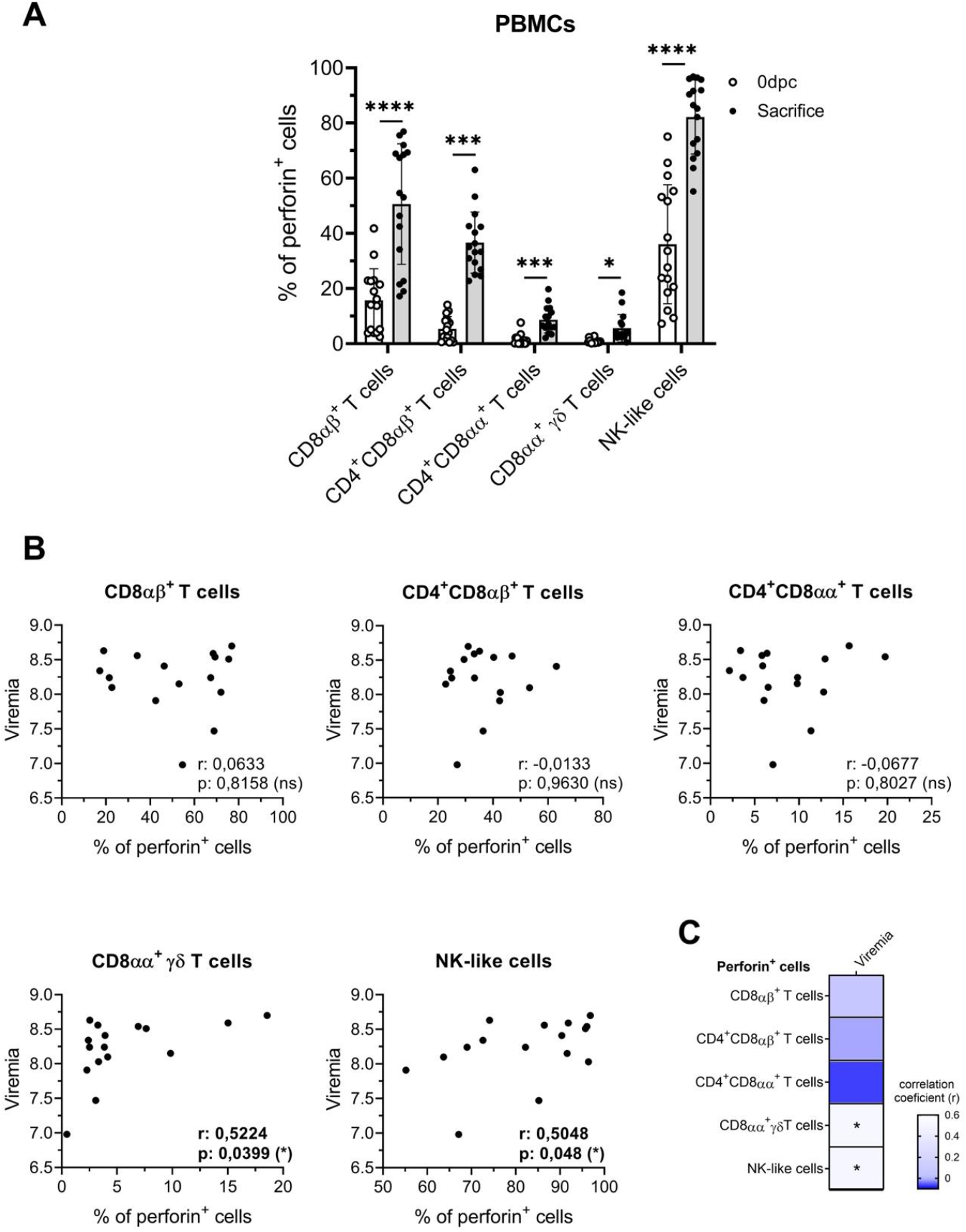
Perforin-producing NK-like and *γ*δ T cells correlate with viral loads during acute ASF. **A)** Frequencies of perforin-producing cytotoxic cell subsets (within the parental populations) measured *ex vivo* in uninfected controls (white bars) and ASFV-infected pigs. **B)** Correlations between frequencies of perforin-producing subsets and blood viral loads during late acute disease. Analyses were performed using cryopreserved PBMCs collected from three independent previous studies. **C)** Heatmap summarising all correlations between frequencies of cytotoxic subsets and viremia. Correlations were assessed using Pearson’s or Spearman’s test (p > 0.05 ns; * p ≤ 0.05; ** p ≤ 0.01; *** p ≤ 0.001; **** p ≤ 0.0001). Significant correlations are highlighted in bold.

Acute ASF is characterised by pronounced lymphopenia during the later stages of infection^50,51^. To determine whether increases in cytotoxic cell frequencies were simply a consequence of global lymphocyte loss, we infected five pigs with Georgia2007/1 and quantified both relative frequencies and absolute counts of blood cell populations at 0 and 7 days post-infection. Total T-cell counts confirmed progressive lymphopenia (**Figure S10A and B**). When perforin-positive and -negative populations were analysed separately, the frequency of perforin-producing cells increased after infection, whereas perforin-negative cells either declined or remained stable (**Figure S10C and D**). Absolute counts further revealed that perforin-producing subsets were largely preserved during infection, and in some cases even expanded, as observed for CD4^+^CD8αα^+^ T cells, in contrast to their perforin-negative counterparts (**Figure 9**). Together, these findings indicate that cytotoxic cells exhibit relative resistance to bystander apoptosis, and may undergo active turnover or partial replenishment during acute ASFV infection despite systemic lymphocyte depletion.

**Figure 9.**
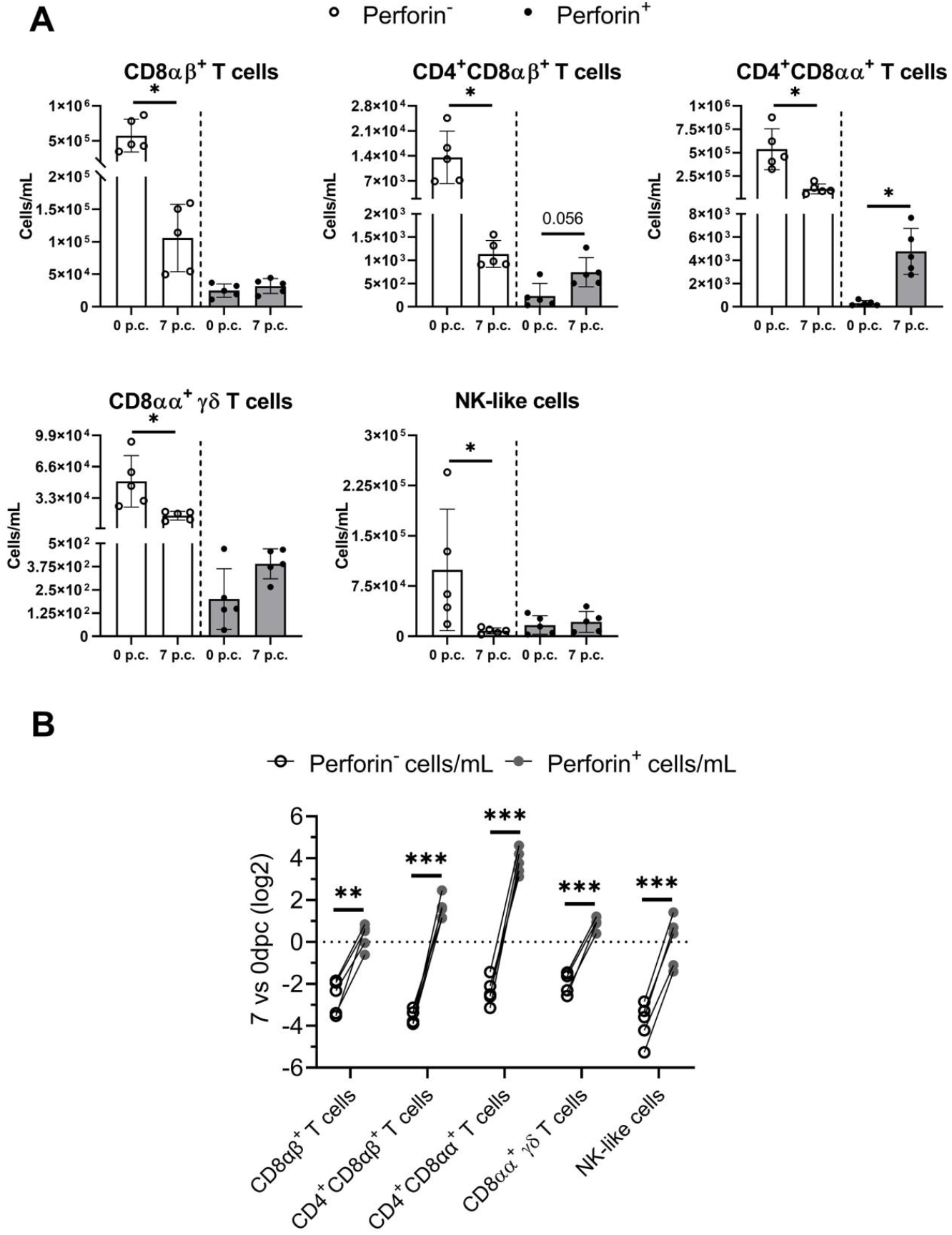
Perforin-producing cytotoxic cells are less susceptible to lymphopenia during acute ASF. **A)** Absolute counts of perforin-positive and perforin-negative populations within the same cytotoxic cell subsets in blood at 0 and 7 days post-infection with Georgia2007/1. Statistical analysis was performed using multiple Wilcoxon tests with Holm-Šidák correction (* p ≤ 0.05). **B)** Fold-change in these populations at day 7 relative to pre-infection levels. Statistical significance was assessed by two-way ANOVA followed by Šidák’s multiple comparisons test (p > 0.05 ns; * p ≤ 0.05; ** p ≤ 0.01; **** p ≤ 0.0001).

## DISCUSSION

Exposure of pigs to either virulent or live attenuated ASFV strains leads to diametrically opposed outcomes, a rapidly fatal disease or the establishment of protective immunity. Although these contrasting trajectories are well characterised, the underlying immune responses remain poorly understood. Here, we show that cytotoxic responses are differentially engaged in these two contexts. During lethal infection with the highly virulent Georgia2007/1 strain, pigs react with a broad cytotoxic response involving several lymphocyte subsets. In contrast, the protective cytotoxic activity induced by the LAV BA71ΔCD2 is primarily mediated by CD8αβ T cells. We also identify additional correlates of protection, underscoring the multifactorial nature of ASF immunity. Together, our data highlight that host response against ASFV depends on complex, multi-layered immune mechanisms that ultimately determine infection outcome.

Vaccine-induced cytotoxic T cells are critical against many viral infections, particularly those in which neutralising antibodies play a limited role, as is the case for ASFV^52,53^. Although this is especially relevant for non-cytopathic viruses^54^, cytotoxic responses also contribute to protection against cytopathic viruses^55–59^. ASFV encodes several early-expressed anti-apoptotic proteins that delay host cell death^17^, suggesting that rapid cytotoxic killing of infected cells may limit viral amplification. Consistent with this idea, vaccine-elicited cytotoxic CD8^+^ T cells might mediate early control of ASFV replication following challenge, as described for other cytopathic viruses^60–63^. The presence of cytotoxic T cells during recall responses in BA71ΔCD2-immunised pigs are in line with previous studies using diverse attenuated ASFV strains^26,29–31,33,64^. However, a detailed characterisation of these responses had been lacking. The depletion experiment by Oura et al.^27^ demonstrated the importance of CD8α^+^ cells in protection, which is expressed in both cytotoxic CD4^+^ and CD4^−^ CD8αβ^+^ T-cell subsets identified in the present study. Notably, depletion of CD8β^+^ cells in one immune pig resulted in lethal infection^27^, supporting the critical role of CD8αβ^+^ T cells in ASFV immunity. Future work will be required to dissect the relative contributions of individual cytotoxic cell subsets in protection. Interestingly, although CD8αβ^+^ T cells were detected in recall responses *in vitro*, their frequencies did not increase *in vivo* in protected pigs at 7 days post-challenge. This suggests that control of intranasal ASFV infection may occur locally at mucosal entry sites, preventing the development of a measurable systemic response. This is consistent with our intranasal infection model, which more closely mimics natural exposure than the intramuscular challenge used in most previous studies^65^. Considering the recognised importance of tissue-resident T cells in antiviral defence^66,67^, future vaccine efficacy studies should adopt challenge models that better reflect natural ASFV infection dynamics.

In contrast to the protective role of vaccine-induced cytotoxic T cells, we observed a broad cytotoxic activity in pigs suffering acute ASF. This aligns with previous studies showing elevated perforin-producing subsets in diseased pigs^26,37,38,41,42^. These findings underline the importance of a timely cytotoxic response. While an early expansion of cytotoxic cells during the initial phase of the infection is beneficial, a late cytotoxic response during uncontrolled viral replication likely reflects dysregulated immunity. The elevated cytotoxic cell frequencies observed in diseased pigs are likely part of a hyperinflammatory response associated with ASF pathology^68,69^. We identified NK and *γ*δ T cells as the subsets whose cytotoxic activity correlated with viremia, suggesting that these innate populations respond directly to virus expansion, whereas cytotoxic αβ T cells are activated more indirectly through inflammation^69^. Although cytotoxic NK and *γ*δ T cells can contribute to antiviral control^70–74^, they may also promote tissue damage through killing either infected or bystander uninfected cells^75–79^. For ASFV, direct exacerbation of disease through macrophage killing by cytotoxic cells is unlikely given the virus’s cytopathic nature. However, they might contribute to ASF-associated leukopenia characteristic of acute ASF, potentially by bystander cell death^41,80^. To note, absolute numbers of perforin-positive cells during acute disease were maintained or increased, in contrast with their perforin-negative counterparts, indicating an active turnover and preservation of cytotoxic capacity. Overall, the increased activity of NK and *γ*δ T cells during acute ASF reflects a compensatory response to uncontrolled viral replication rather than a direct driver of immunopathology.

Beyond cytotoxic T cells, our results highlight that BA71ΔCD2-induced protection is multifactorial. Similar observations have been made in pigs immunised with the partially attenuated Estonia 2014 strain, which also showed diverse correlates of protection^81^. These findings support the broader concept that effective antiviral immunity requires coordinated engagement of innate, humoral and cellular compartments^82–86^. Synergy between vaccine-induced humoral and cellular responses may reduce the immune threshold required for protection^87^. However, additional mechanistic studies are needed to discriminate true causal correlates from surrogate markers of protection in ASF^88^. Antibodies represent another key component of ASFV immunity^23^. We found a strong correlation between ASFV-specific IgG titres and protection, consistent with observations in pigs immunised with the attenuated Pret4Δ9GL strain^89^. ASFV-specific IgG appears concurrently with the onset of BA71ΔCD2-mediated protection^40^. Although antibody-mediated protection remains incompletely understood, neutralising antibodies induced by ASFV-G-Δ9GL/ΔUK or ASFV-G-ΔI177L LAVs have been associated with protection^90^. However, as these assays were performed with cell-adapted isolates, their physiological relevance remains unclear. The potential role of non-neutralising antibodies may also contribute to ASFV immunity^91–94^, especially given their importance as correlates of protection in other viral systems^95^. Finally, IFN*γ*-producing cells emerged among the multiple correlates of protection, and strongly correlated with the inflammatory transcriptional signature observed in stimulated cells. We previously showed that recall responses in BA71ΔCD2-vaccinated pigs involve a positive feedback loop in which IFN*γ* secreted by Th1 memory cells activates inflammatory macrophages^31^. This rapid induction of innate immunity may restrict ASFV replication in activated macrophages^43,96,97^, contributing to early viral control.

In summary, this study advances the mechanistic understanding of cytotoxic responses in ASFV immunity. We identify CD8αβ T cells as key mediators of vaccine-induced protection and contrast their focused activation with the broad, dysregulated cytotoxic landscape observed during acute disease. Our findings emphasize that protective immunity likely relies on rapid, localised control of viral replication at mucosal entry sites, coupled with coordinated engagement of cytotoxic T cells, antibodies, and IFN*γ*-producing memory cells. These insights contribute to the rational design of safer and more effective ASFV vaccines.

## Acknowledgement

This work was funded by the Spanish Ministry of Science and Innovation, MICIU/AEI/10.13039/501100011033, grant PID2022-136312OB-I00 (F.R. & J.A.). We acknowledge the support of Red de Investigación en Sanidad Animal (RISA) and World Organisation for Animal Health (WOAH). We thank the Animal Facility unit from IRTA-CReSA for their excellent technical support. The Georgia2007/01 ASFV strain was kindly provided by Dr. Linda Dixon, from the Pirbright Institute (UK). All content and code were reviewed and verified by the authors.

