## Supplemental figures for "Distinct cytotoxic cell subsets underlie protective and non-protective immunity to African swine fever virus"

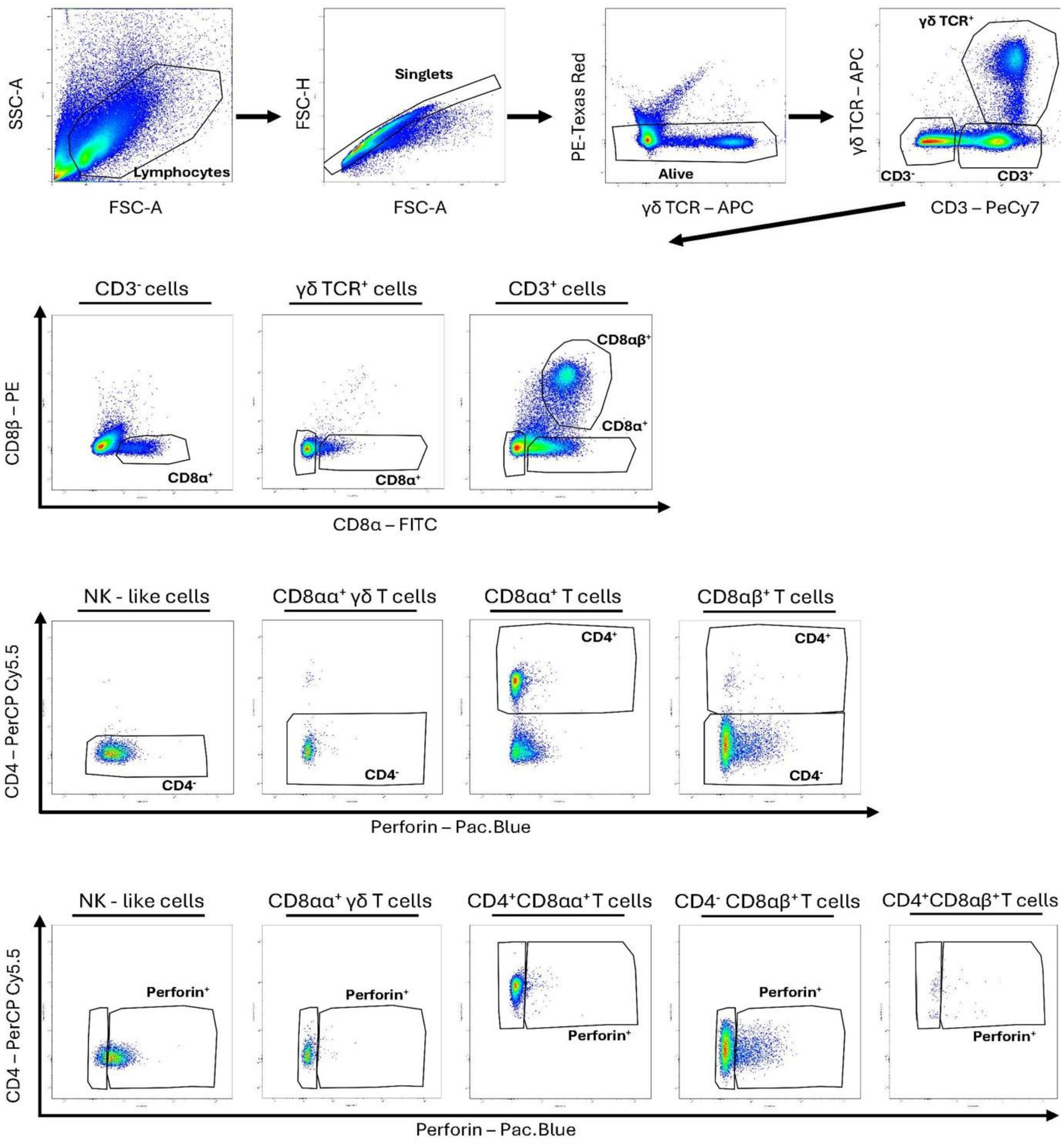


**Figure S1. Flow cytometry gating strategy.** Representative plots used to identify perforin-producing lymphoid cell subsets.

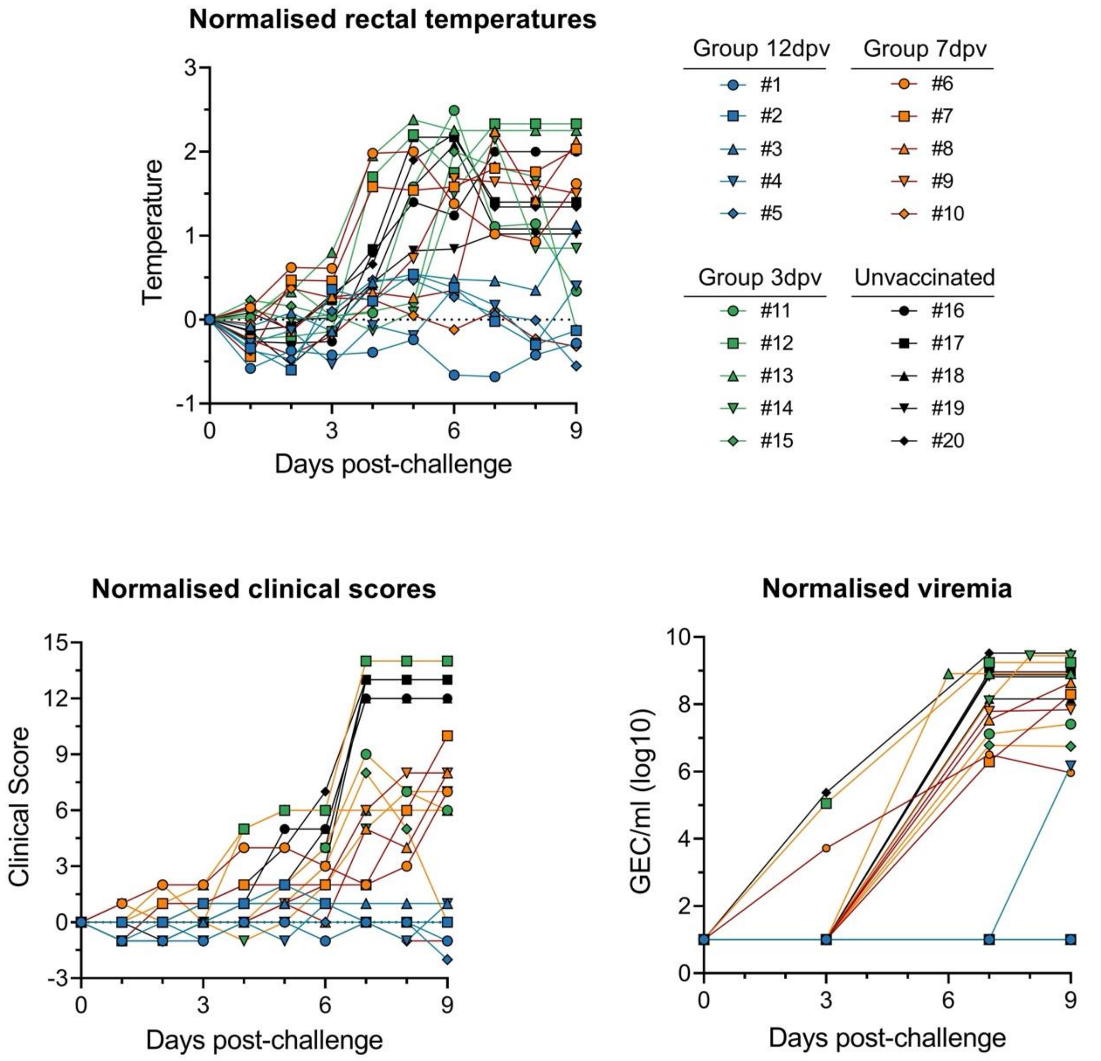


**Figure S2. Kinetics of fever, clinical signs and virus loads after challenge.** Rectal temperatures, clinical scores and viremias from pigs vaccinated 3 (green), 7 (orange) or 12 (blue) days before the challenge. Unvaccinated controls are shown in black. Data were normalised to day 0 post-challenge, and the area under the curve (AUC) were used for correlation analyses.

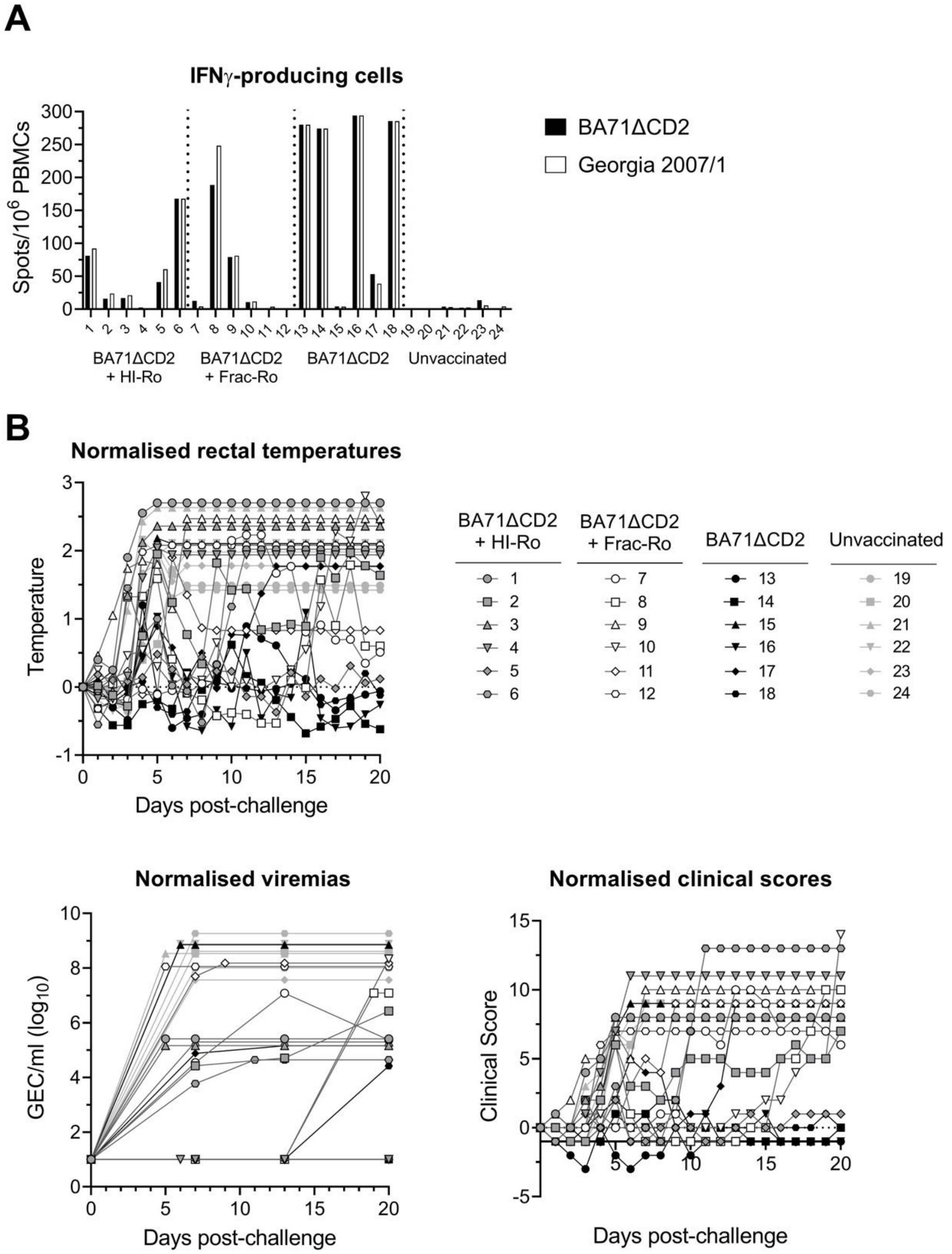


**Figure S3. Normalised rectal temperature, viremia, and clinical scores after challenge. A)** Percentages of IFNγ-secreting PBMCs measured by ELISpot at day 0 post-challenge after stimulation with BA71ΔCD2 (black bars) or Georgia2007/1 (white bars). **B)** Rectal temperatures, clinical scores, and viremias from pigs vaccinated with BA71ΔCD2 alone or with HI-Ro or Frac-Ro; unvaccinated controls are included. Data were normalised to day 0 post-challenge, and the area under the curve (AUC) values were used for correlation analyses. GEC: genomic equivalent copies.

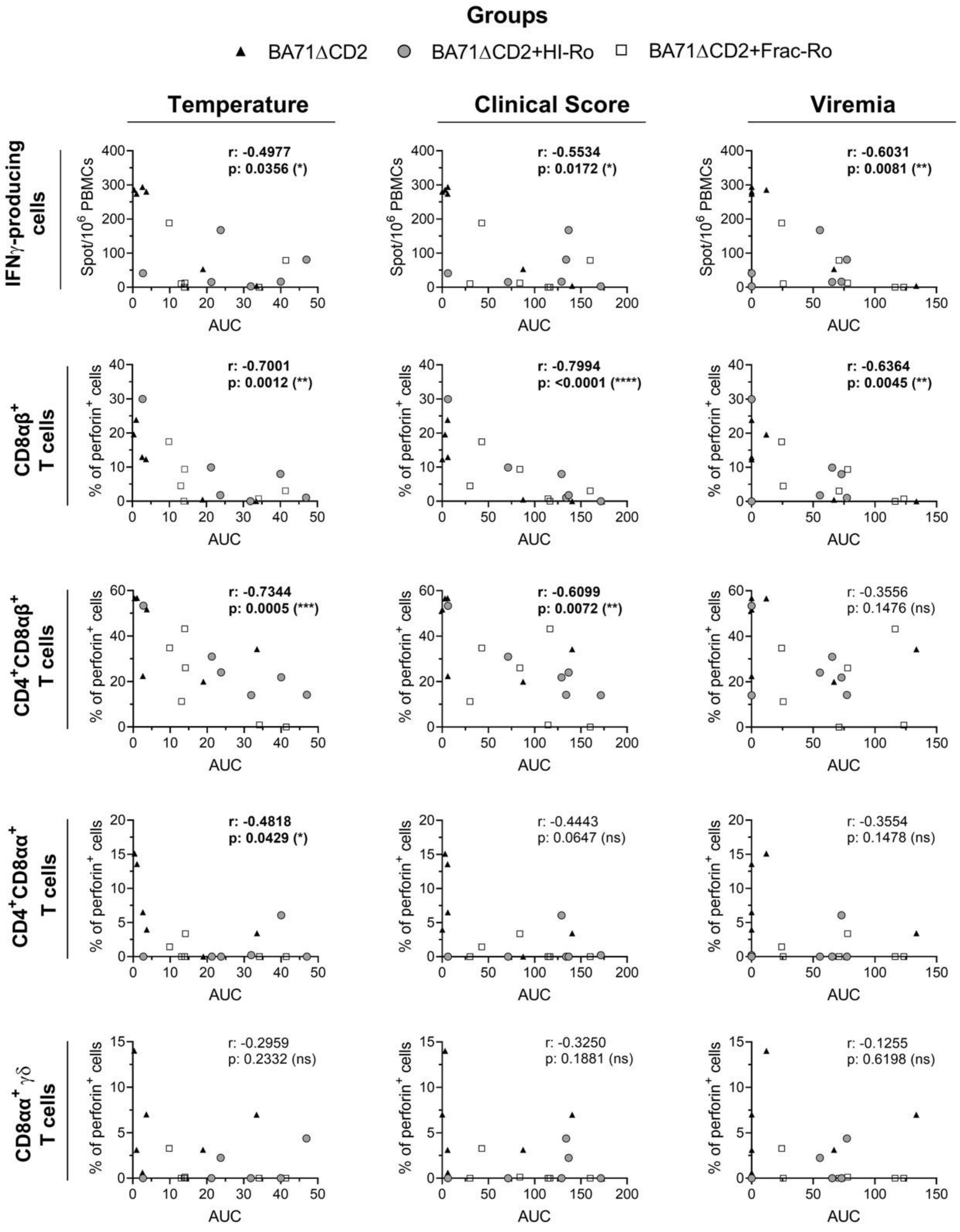


**Figure S4. Correlations between disease control, IFNγ-secreting cells, and cytotoxic cells after BA71ΔCD2 stimulation.** Frequencies of IFNγ-secreting and perforin-producing PBMCs (within the parental populations) before challenge were quantified after stimulation with BA71ΔCD2 and correlated with normalised AUC values of rectal temperature, clinical scores, or viremia. Circles: BA71ΔCD2+HI-Ro; squares pigs: BA71ΔCD2+Frac-Ro; triangles: BA71ΔCD2 alone. Correlations were assessed using Pearson's or Spearman's test, as appropriate ($p > 0.05$ ns; * $p \leq 0.05$; ** $p \leq 0.01$; *** $p \leq 0.001$; **** $p \leq 0.0001$). Significant correlations are highlighted in bold.

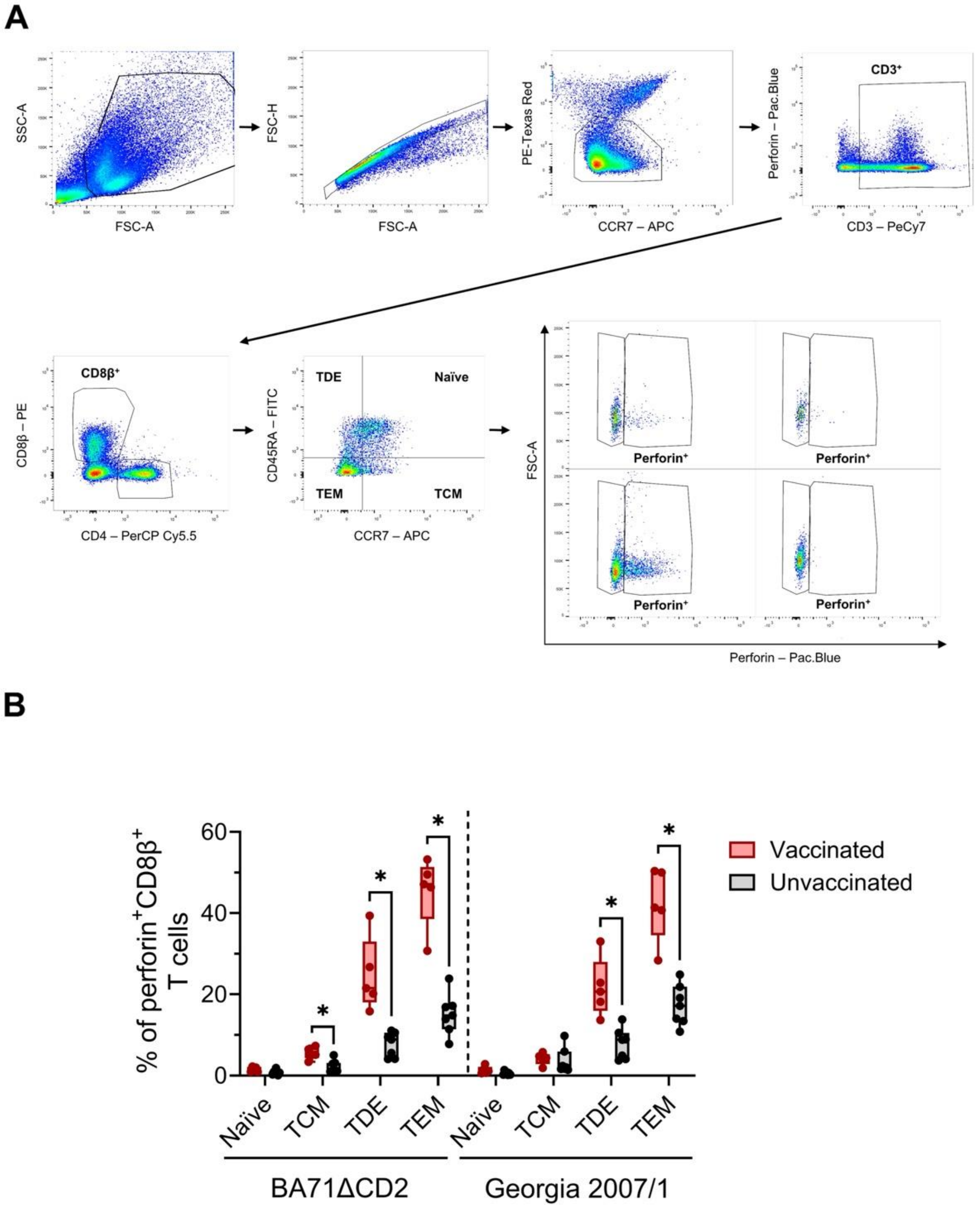


**Figure S5. Immunophenotyping of BA71ΔC2-induced cytotoxic CD8β$^{+}$ T cells. A)** Flow cytometry gating strategy used to classify perforin$^{+}$ CD8β$^{+}$ T cells into naïve (CD45RA$^{+}$ CCR7$^{+}$), central memory (TCM; CD45RA$^{-}$ CCR7$^{+}$), effector memory (TEM; CD45RA$^{-}$ CCR7$^{-}$), and terminally differentiated effector (TDE; CD45RA$^{+}$ CCR7$^{-}$) subsets. **B)** PBMCs from BA71ΔCD2-vaccinated pigs showing strong protection were stimulated with BA71ΔCD2 or Georgia2007/1 to assess perforin-producing CD8β$^{+}$ T-cell phenotypes. Statistical analyses were performed using multiple Mann-Whitney tests with Holm-Šidák correction (* $p \leq 0.05$).

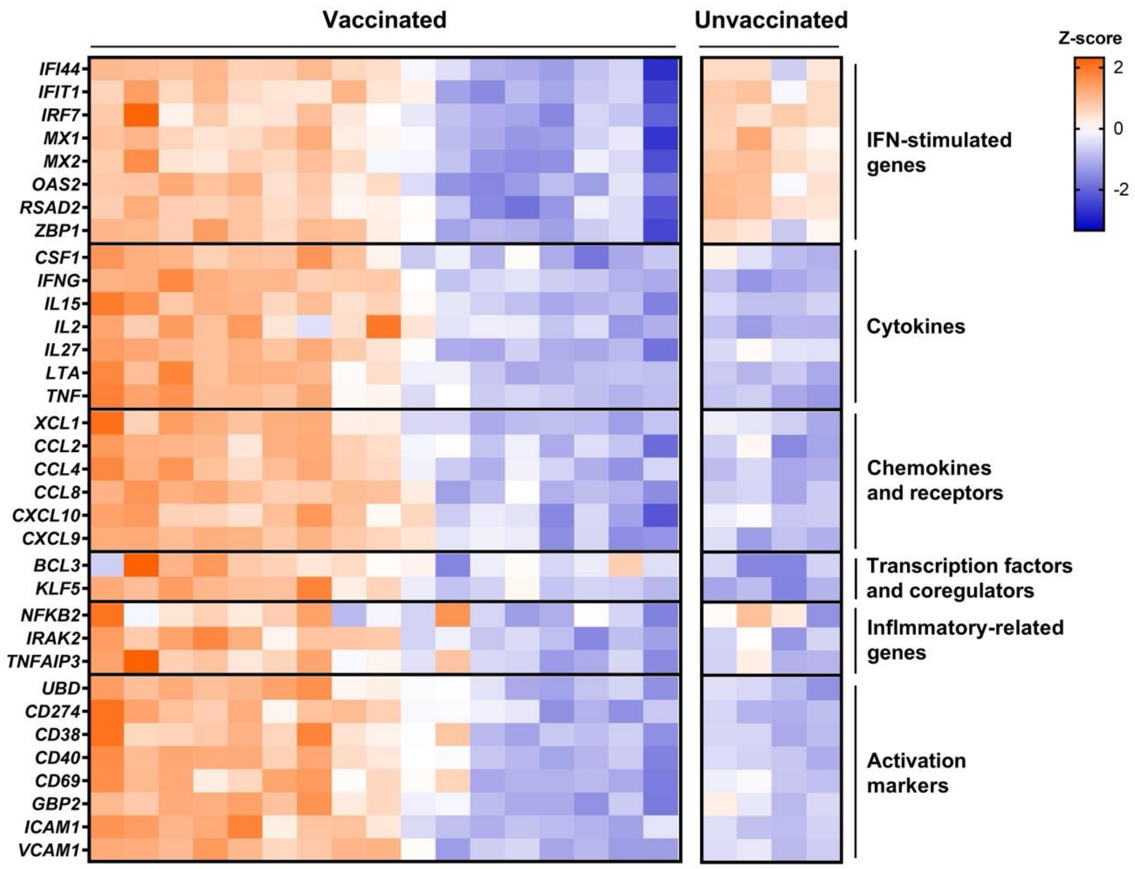


**Figure S6. Innate immune transcriptomic signatures after BA71∆CD2 stimulation.** PBMCs collected at day 0 post-challenge were stimulated with BA71∆CD2. The heatmap shows z-score normalised $\log_2$ fold change ($\log_2$FC) values for innate immune-related genes compared with unstimulated cells.

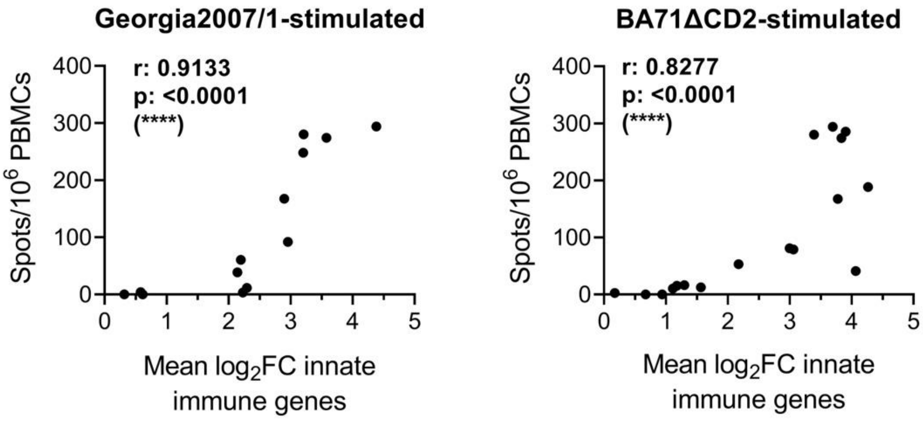


**Figure S7. IFNγ-secreting cells correlate strongly with innate immune signatures during the recall response**. Correlation analyses between IFNγ-secreting PBMCs (ELISpot) and mean of the $\log_2$ fold change ($\log_2$FC) values of innate immune genes after stimulation with Georgia2007/1 or BA71ΔCD2. Correlations were assessed using Pearson's or Spearman's test (**** $p \leq 0.0001$).

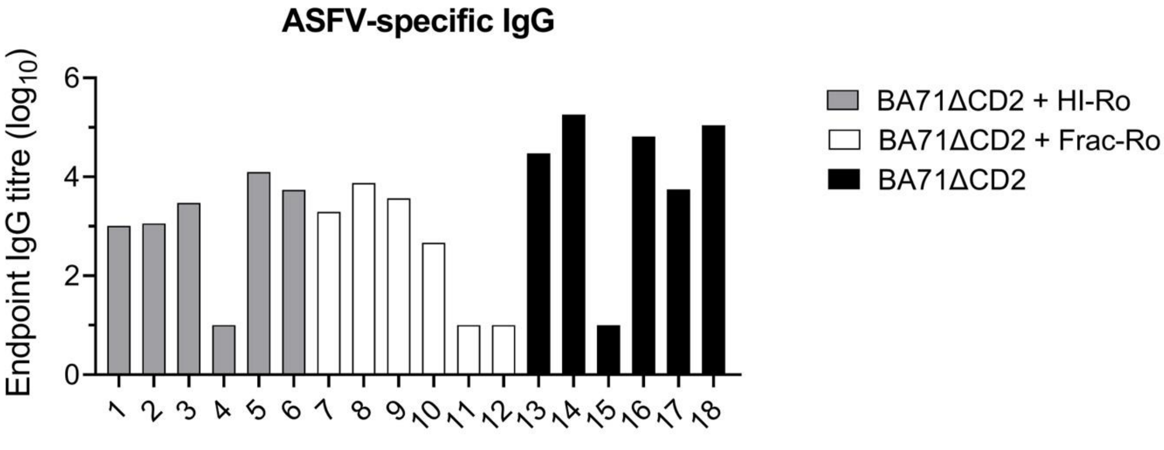


**Figure S8. ASFV-specific IgG endpoint titres at day 0 post-challenge.** IgG titres from pigs vaccinated with BA71ΔCD2 alone or in combination with HI-Ro or Frac-Ro.

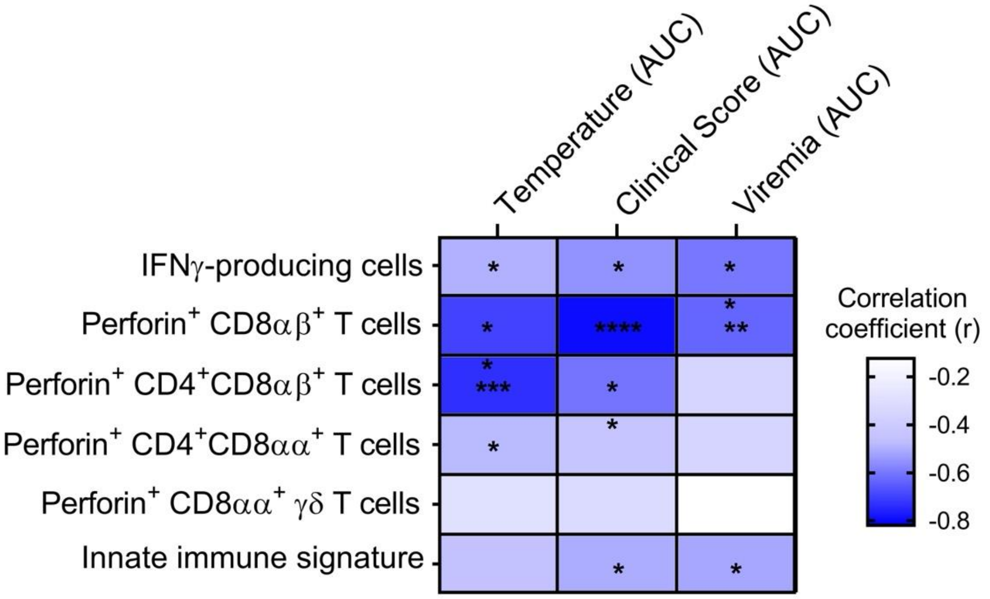


**Figure S9. BA71ΔCD2-specific correlates of protection**. Heatmap summarising all correlations between ASFV-specific immune responses measured upon BA71ΔCD2 stimulation and ASF disease control. Correlations were assessed using Pearson's or Spearman's test ($p > 0.05$ ns; * $p \leq 0.05$; ** $p \leq 0.01$; *** $p \leq 0.001$; **** $p \leq 0.0001$).

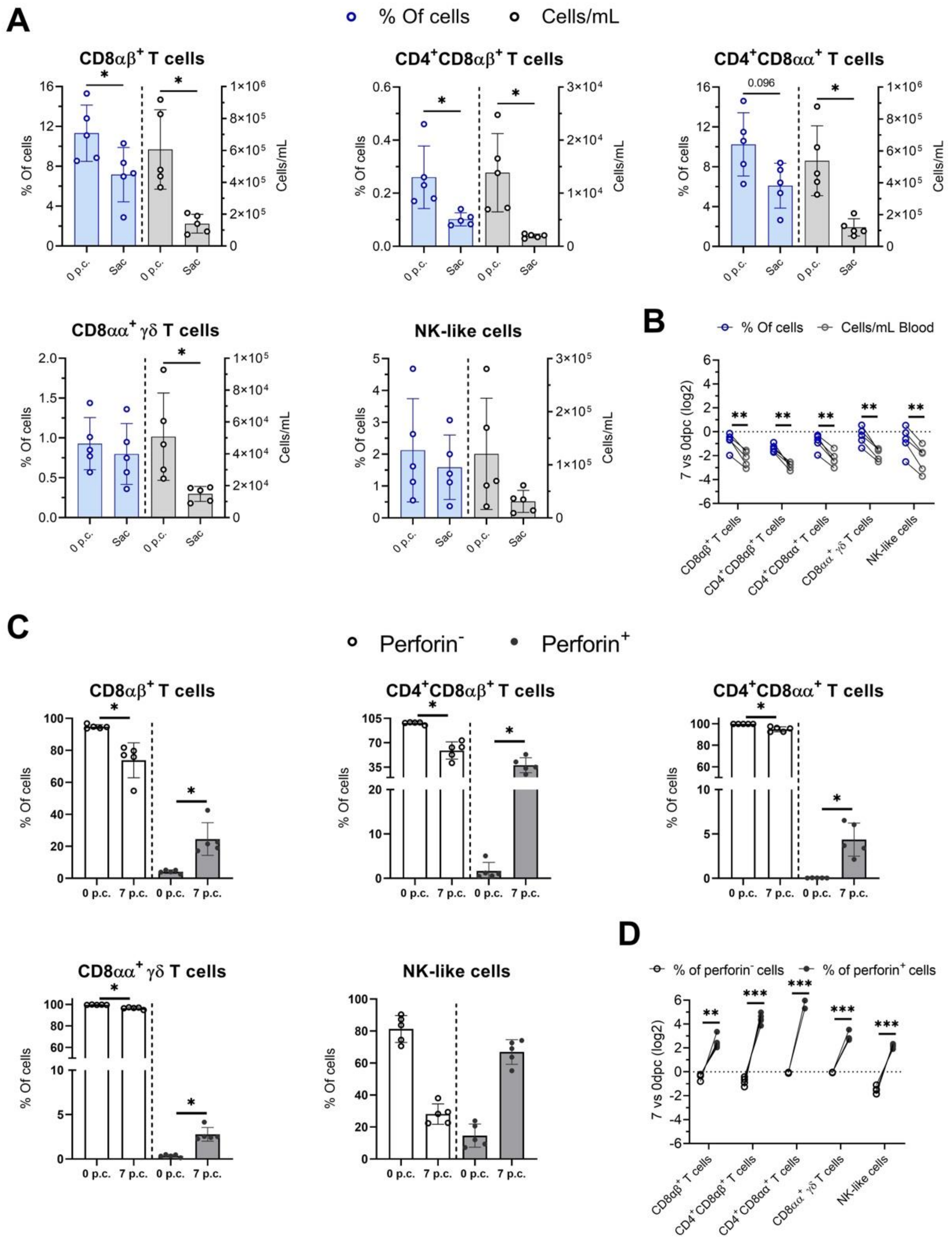

**Figure S10. Increased frequencies of cytotoxic subsets during acute ASF-associated lymphopenia**. **A)** Frequencies (within live PBMCs) and absolute counts of the indicated cell populations in blood at 0 and 7 days post-infection with Georgia2007/1. **B)** Fold-change in these populations at day 7 relative to pre-infection levels. **C)** Frequencies of perforin-positive and perforin-negative populations (within parental subsets) in blood at 0 and 7 days post-infection. **D)** Fold-change of these perforin-defined populations at day 7 relative to pre-infection levels. Statistical analyses for panels A and C were performed using multiple Wilcoxon tests with Holm-Šidák correction (* $p \leq 0.05$). Significance for panels B and D was assessed by two-way ANOVA followed by Šidák's multiple comparisons test ($p > 0.05$ ns; * $p \leq 0.05$; ** $p \leq 0.01$; **** $p \leq 0.0001$).
